## Supplemental file for "Loss of Bone Morphogenetic Protein-binding Endothelial Regulator Causes Insulin Resistance"

**Supplementary Table 1. Metabolic parameters.**

| Parameters | WT | iKO |
|---|---|---|
| BW (g) | 32.25 ± 0.86 | 32.18 ± 1.05$^{(NS)}$ |
| Glucose (mg/dl) | | |
| Fasted | 139.0 ± 4.79 | 150.0 ± 3.81$^{(NS)}$ |
| Fed | 174.7 ± 6.12 | 176.4 ± 8.134$^{(NS)}$ |
| Insulin (ng/ml) | | |
| Fasted | 1.31 ± 0.13 | 2.80 ± 0.22$^{c}$ |
| Fed | 2.81 ± 0.20 | 16.77 ± 0.42$^{d}$ |
| TG (mg/dl) | 73.12 ± 4.25 | 85.78 ± 3.30$^{a}$ |
| FFA (mEq/I) | 0.78 ± 0.10 | 0.78 ± 0.05$^{(NS)}$ |

NS, not significant.

$^{a}$ P<0.05, $^{b}$ P<0.01, $^{c}$ P<0.001, $^{d}$ P<0.0001

WT, BMPER$^{flox/flox}$;CAG-CreER$^{-/-}$.

iKO, BMPER$^{flox/flox}$;CAG-CreER$^{+/-}$.

**Supplementary Table 2. Mass spectrometry result shows representative proteins that potentially form complex with BMPER.**

| Gene ID | GeneSymbol | SRA | PSMs | iBAQ |
|---|---|---|---|---|
| 3939 | LDHA | S | 118 | 0.38 |
| 4000 | LMNA | S | 100 | 0.17 |
| 8514 | KCNAB2 | S | 49.2 | 0.12 |
| 4864 | NPC1 | S | 19.2 | 0.05 |
| 54887 | UHRF1BP1 | S | 13.5 | 0.02 |

S: high confidence; R: medium confidence; A: weak confidence

PSMs: total normalized peptide hits

iBAQ: total abundance of the gene

**Supplementary Table 3. Main reagent table.**

| REAGENT or RESOURCE | SOURCE | IDENTIFIER |
|---|---|---|
| **Antibodies** | | |
| LRP1-CTD | Sigma | Cat#L2170 |
| BMPER (Crossveinless-2/CV-2) | R&D Systems | Cat# AF2299 |
| NPC1 | Abcam | Cat# 124801 |
| IRS-1 | Cell signaling | Cat#2382 |
| phospho- IRS1 (Tyr612) | Millipore | Cat#09-432 |
| Akt | Cell signaling | Cat#9272 |
| phospho-Akt (Ser473) | Cell signaling | Cat#9271 |
| phospho-Akt (Thr308) (D25E6) | Cell signaling | Cat#13038 |
| Insulin Receptor β (4B8) | Cell signaling | Cat#3025 |
| β-Actin (C4) -HRP | Santa Cruz | Cat#sc-47778 |
| Smad1 | Cell signaling | Cat#9743 |
| phospho- Smad1/Smad5/Smad8 (Ser463/465) | EMD Millipore | Cat#AB3848-I |
| Flag antibody | Sigma-Aldrich | Cat#2426 |
| EZview Red ANTI-FLAG M2 Affinity Gel | Sigma-Aldrich | Cat#F2426 |
| **Chemicals, Peptides, and Recombinant Proteins** | | |
| 60% high-fat diet (HFD) | Research Diets | Cat#D12492 |
| Insulin | Novolin R | Cat#0169-1833-11 |
| Mouse BMPER (Crossveinless-2) Protein | R&D Systems | Cat#2299-CV-050 |
| Protein A/G Plus-agarose beads | Santa Cruz | Cat#sc-2003 |
| iScript$^{TM}$ cDNA synthesis kit | Bio-Rad Laboratories | Cat#1708891 |

| | | |
|---|---|---|
| iTaq SYBR Green supermix | Bio-Rad Laboratories | Cat#1725121 |
| TaqMan™ Universal PCR Master Mix | Thermo Scientific | Cat#4304437 |
| Insulin solution | Santa Cruz | Cat# 11061-68-0 |
| Cell maintenance supplements | Gibco | Cat# A15564 |
| Tamoxifen | Sigma-Aldrich | Cat#T5648 |
| Lipofectamine 2000 | Thermo Scientific | Cat#11668019 |
| Perfusion Medium | Thermo Scientific | Cat#17701038 |
| Collagenase | Sigma-Aldrich | Cat#C5168 |
| DMEM | Thermo Scientific | Cat#11965118 |
| William's E Medium | Thermo Scientific | Cat#12551-032 |
| Immobilon-P transfer membrane | Millipore | IPVH00010 |
| Wash Medium | Thermo Scientific | Cat#17704-024 |
| Percoll | GE Healthcare | Cat#17089101 |
| OptiPrep density gradient medium | Sigma-Aldrich | Cat#1556 |
| Penicillin & Streptomycin | Thermo Scientific | Cat#15140122 |
| Fetal bovine serum (FBS) | Sigma-Aldrich | Cat#F0926 |
| Horse serum | Thermo Scientific | Cat#16050114 |
| Maintenance supplements | Thermo Scientific | Cat#CM4000 |
| Chlorpromazine (CPM) | Sigma-Aldrich | Cat#C8138 |
| Methyl-β-cyclodextrin (MCD) | Sigma-Aldrich | Cat#M7439 |
| **Critical Commercial Assays** | | |
| Insulin ELISA kit | Millipore | Cat#EZRMI-13K |
| RNA purification kit | QIAGEN | Cat#74104 |
| Infinity Glucose kit | Thermo Scientific | Cat#TR15421 |
| Albumin ELISA kit | Abcam | Cat#ab108792 |

| | | |
|---|---|---|
| **Experimental Models: Cell Lines** | | |
| Primary hepatocytes | This paper | N/A |
| HEK293 cells | ATCC | Cat#CRL-1573 |
| **Experimental Models: Organisms/Strains** | | |
| Mouse: B6.Cg-Tg (CAG-cre/ Esr1*) 5Amc/J (CAG-CreER$^{+/-}$) | Jackson Laboratories | JAX: 004682 |
| Mouse: Cdh5(PAC)-CreERT2 (Cdh5-CreER$^{+/-}$) | Dr. Ralf H. Adams from Max Planck Institute for Molecular Biomedicine, Germany | N/A |
| Mouse: C57BL/6 | Jackson Laboratories | JAX: 000664 |
| Mouse: IR$^{flox}$ | Jackson Laboratories | JAX: 006955 |
| Mouse: B6 db | Jackson Laboratories | JAX: 000697 |
| **Recombinant DNA** | | |
| Npc1 mouse shRNA plasmid | Origene | Cat#TL501501 |
| pCMV-Tag2 vector | Agilent | Cat#211172 |
| Scrambled shRNA plasmid | OriGene | Cat#TR30021 |
| pLenti-C-Flag-DDK-BMPER | This paper | N/A |
| Ad-CMV-iCre | Vector Biolabs | Cat#1045 |

**Supplementary Table 4. Sequence information for qPCR primers.**

| **Gene** | Forward primer | Reverse primer |
|---|---|---|
| G6Pase | 5'- tctgtcccggatctaccttg-3' | 5'- gaaagtttcagccacagcaa-3' |
| PEPCK | 5'- ggagtacccattgagggtatcat-3' | 5'- gctgagggcttcatagacaag-3' |
| GK | 5'- cagatgctggatgacagagc-3' | 5'- gccaggatctgctctacctttt-3' |
| SREBP1 | 5'- acaagattgtggagctcaaagac-3' | 5'- tgcgcaagacagcagattta-3' |
| IL1β | 5'- agttgacggaccccaaaag-3' | 5'- agctggatgctctcatcagg-3' |
| IL6 | 5'- gctaccaaactggatataatcagga-3' | 5'- ccaggtagctatggtactccagaa-3' |
| TNFα | 5'- ctgtagcccacgtcgtagc-3' | 5'- ttgagatccatgccgttg-3' |
| β-Actin | 5'- ccaaccgtgaaaagatgacc-3' | 5'- accagaggcatacagggaca-3' |
| BMPER | 5'- ggctgagccatgtgtcct-3' | 5'- cgcacctcagactctgtcac-3' |
| BMP2 | 5'- agatctgtaccgcaggcact-3' | 5'- gttcctccacggcttcttc-3' |
| BMP4 | 5'- gatctttaccggctccagtct-3' | 5'- tgggatgttctccagatgttc-3' |
| BMP6 | 5'- actgactagcgcgcagga-3' | 5'- tgtggggagaactccttgtc-3' |
| BMP7 | 5'- cgagaccttccagatcacagt-3' | 5'- cagcaagaagaggtccgact-3' |
| BMP9 | 5'- ggaagctgtgggtagatgacc-3' | 5'- caagtcggtggggatgat-3' |
| BMPR2 | 5'- gagccctcccttgacctg-3' | 5'- gtatcgaccccgtccaatc-3' |

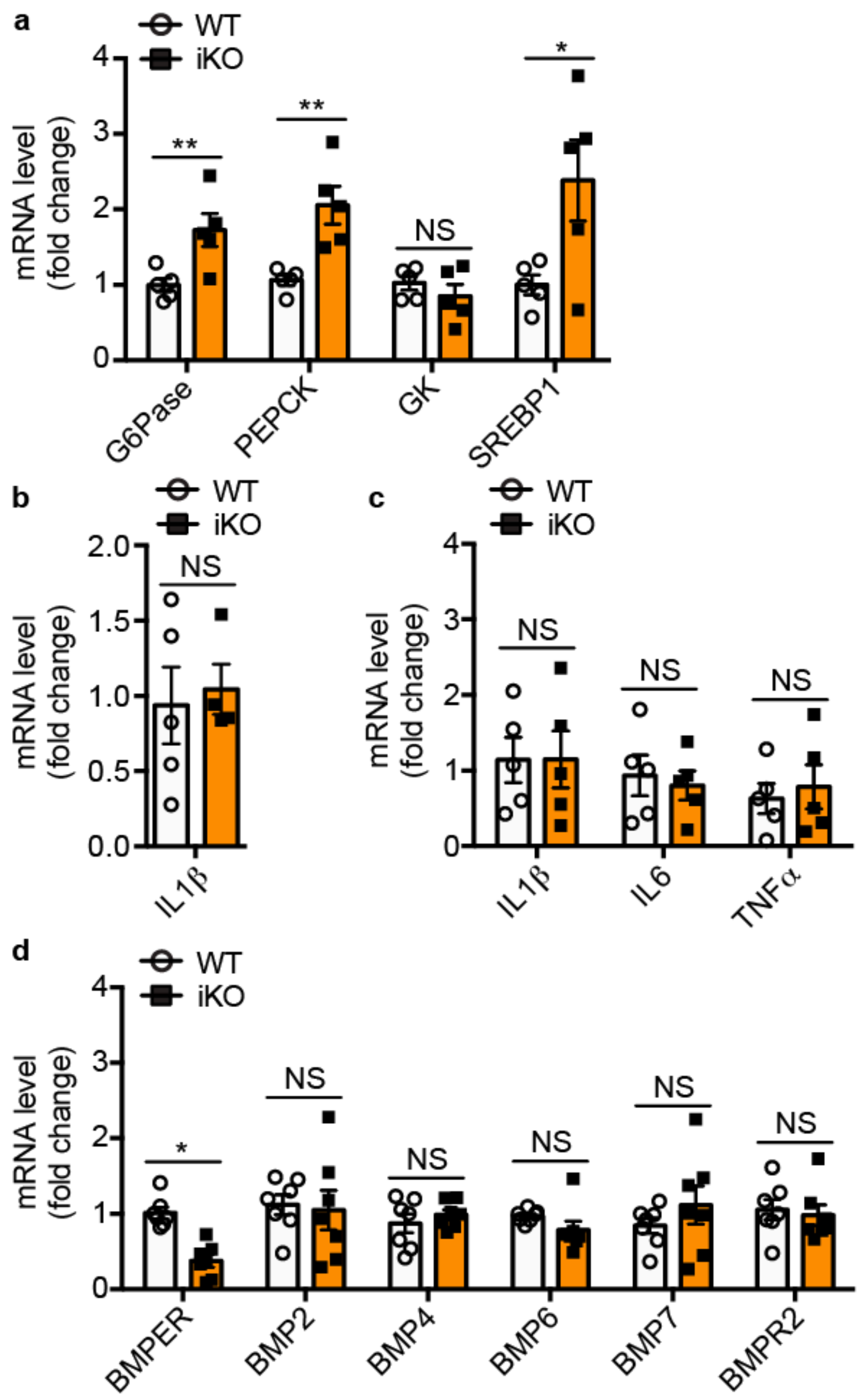


**Supplementary Figure 1. Gene expression changes in liver and WAT tissues of BMPER iKO mice.** (**a-d**) Real-time PCR assays were performed for indicated genes in the liver (**a, b, d**) and WAT (**c**) of BMPER iKO and their littermate control (WT) mice. n=5 (**a-c**) and 7 (**d**). *, *P*< 0.05. **, *P*< 0.01. NS, not significant.

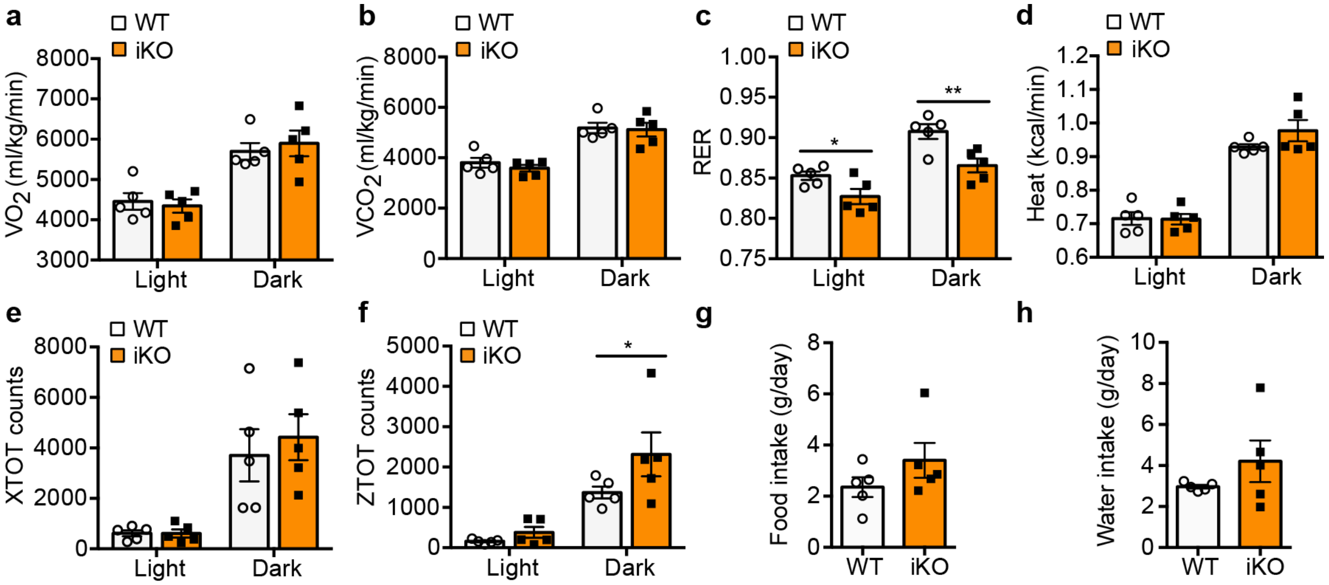


**Supplementary Figure 2. CLAMS studies with BMPER iKO mice.** Indirect calorimetry studies were performed to evaluate $VO_2$ (**a**), $VCO_2$ (**b**), respiratory exchange ratio (RER, **c**), heat (**d**), locomotor activity in x axis (XTOT, **e**) and z axis (ZTOT, **f**), daily food (**g**) and water intake (**h**). n=5 (**a-h**). *, $P < 0.05$. **, $P < 0.01$.

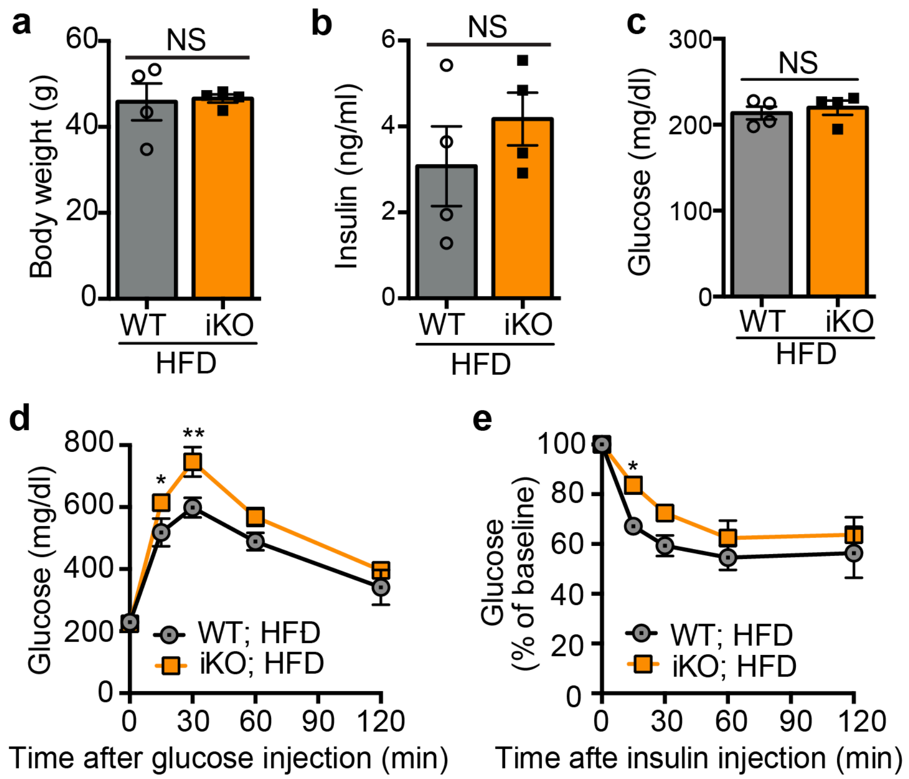


**Supplementary Figure 3. BMPER depletion exacerbates glucose responses in HFD-fed mice.** Metabolic studies were performed with BMPER iKO and WT mice following eight weeks of HFD feeding. (**a**) Body weight. (**b-c**) Fasted insulin and glucose. (**d-e**) Glucose and insulin tolerance tests. HFD, high-fat diet. n= 4 (**a-e**). *, *P*< 0.05. **, *P*< 0.01. NS, not significant.

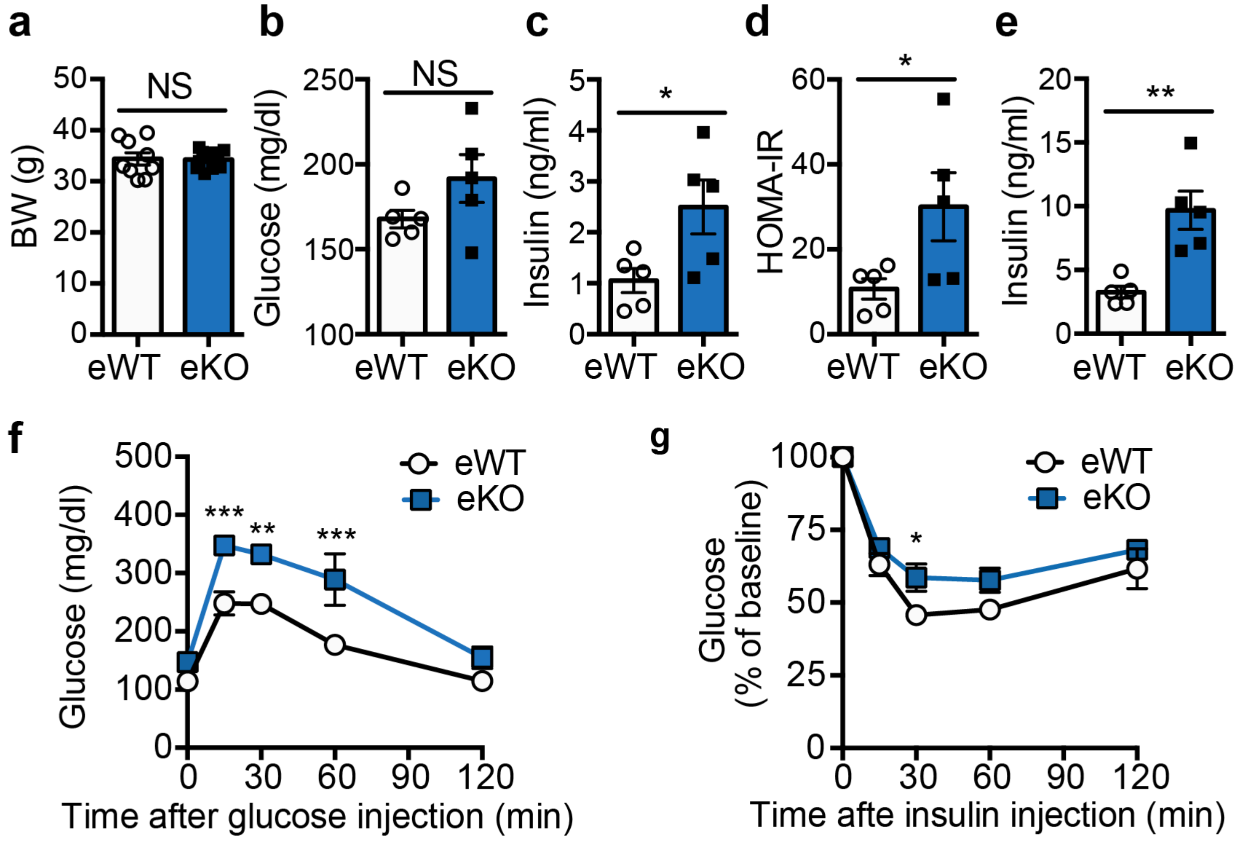


**Supplementary Figure 4. BMPER depletion in ECs leads to glucose dysregulation.** Metabolic studies were performed with BMPER eKO and eWT mice at four months after tamoxifen injection. (**a**) Body weight (BW). (**b-d**) Fasted glucose and insulin, HOMA-IR. (**e**) Fed insulin. (**f-g**) Glucose and insulin tolerance tests. eWT, BMPER$^{flox/flox}$; Cdh5-CreER$^{-/-}$. eKO, BMPER$^{flox/flox}$; Cdh5-Cre ER$^{+/-}$. n=5 (**a-g**). *, *P*< 0.05. **, *P*< 0.01. ***, *P*< 0.001. NS, not significant.

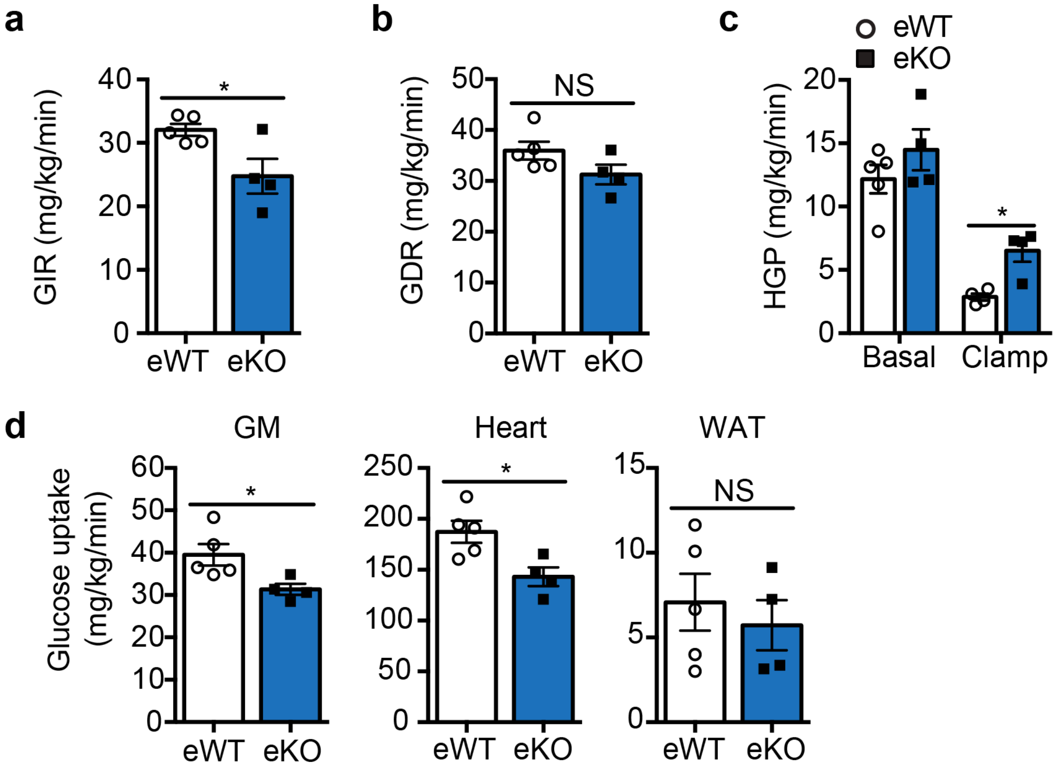


**Supplementary Figure 5. BMPER depletion in ECs leads to insulin resistance.** Clamp studies were performed with BMPER eKO and eWT mice at four months after tamoxifen injection. (**a**) Glucose infusion rate (GIR). (**b**) Glucose disposal rate (GDR). (**c-d**) Hepatic glucose production (HGP; **c**) and glucose uptake in peripheral tissues (**d**) were analyzed with hyperinsulinemic-euglycemic clamps. GM, gastrocnemius muscle; WAT, white adipose tissue. n=5 (**a-d**). *, $P< 0.05$. NS, not significant.

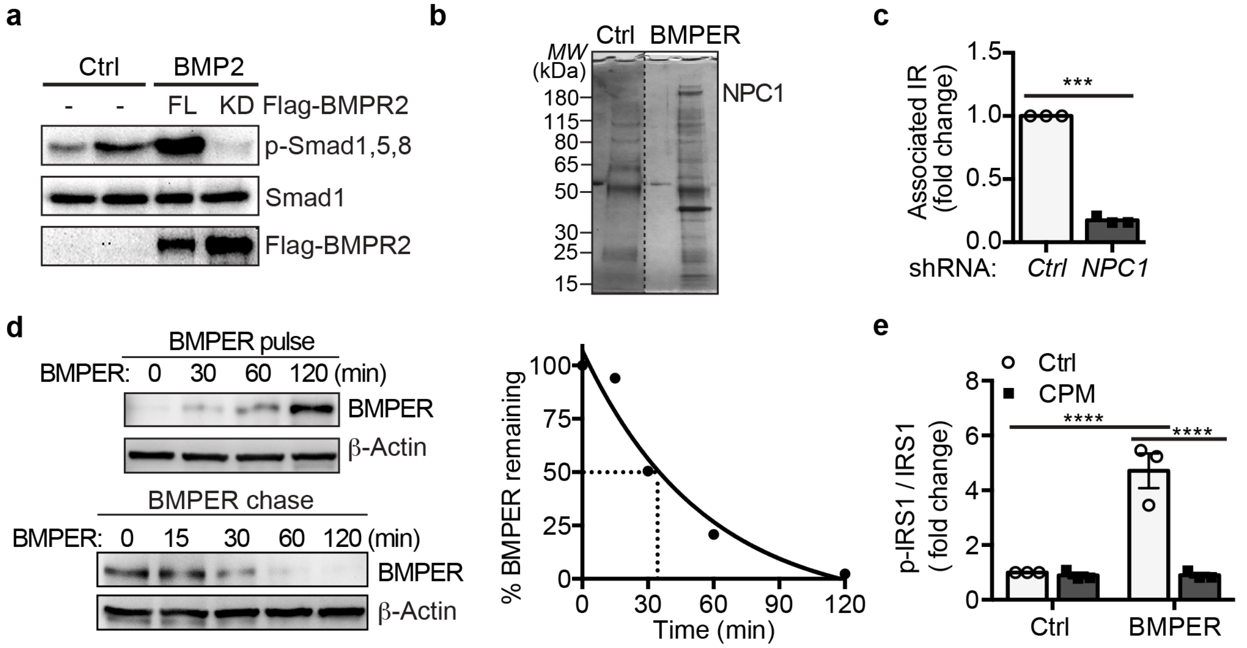


**Supplementary Figure 6. The regulation of BMPER-promoted insulin signaling pathway.** Hepatocytes were transduced with adenovirus of flag-tagged BMPR2 full length (FL) or kinase-dead mutant (KD). Cells were then treated with BMP2 (4 nM, 1 hr). (**b**) NPC1 was identified in BMPER-bound complex. HEK293 cells were treated with BMPER (30 nM) and lysates were immunoprecipitated with anti-BMPER antibody and stained with Coomassie blue. Selected bands were subjected for mass spectrometry analysis. (**c**) Quantitative data for Figure 4e. (**d**) Hepatocytes were pulsed with Flag-BMPER (100 nM, 7.2 ng) for indicated time periods (left top panel), and then chased at 4°C (left bottom panel). Right panel is graphic representation of normalized intracellular BMPER level from the cold chase. (**e**) Quantitative data for Figure 4h. n=3 (**c, e**). ***, *P*< 0.001. ****, *P*< 0.0001.

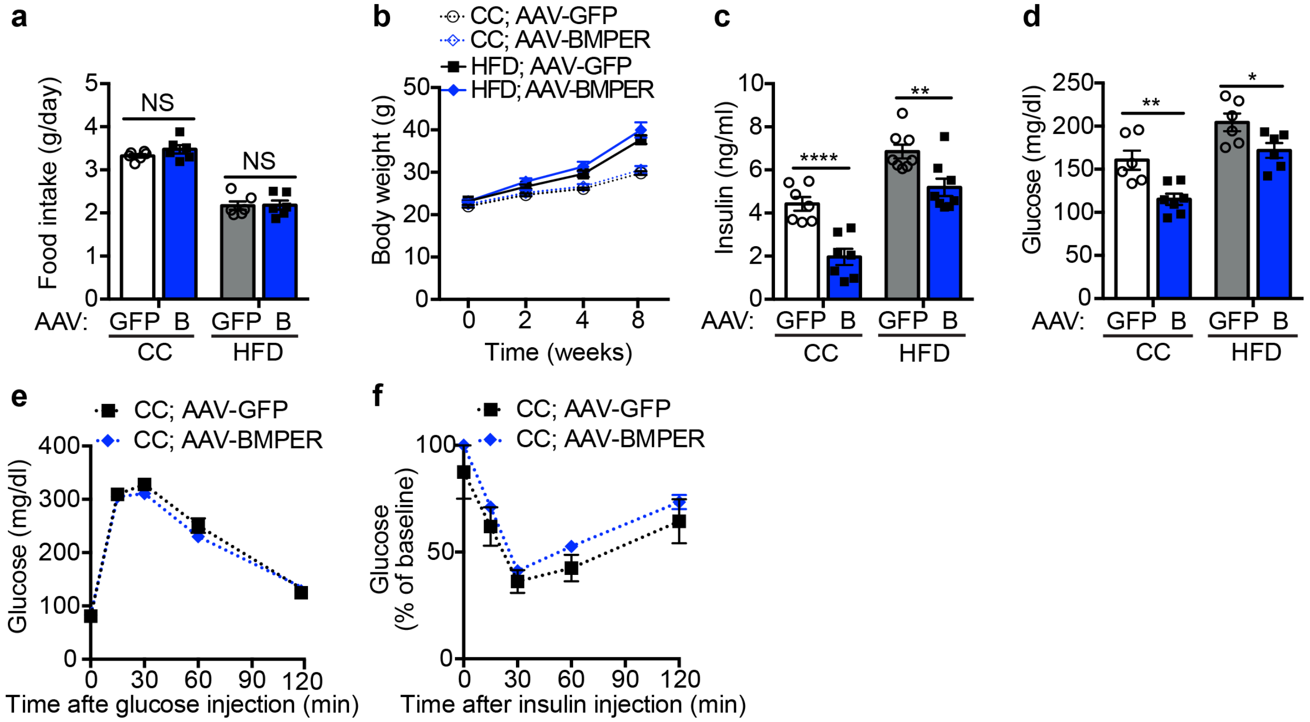


**Supplementary Figure 7. AAV-BMPER improves glucose responses in DIO mice.** AAV-BMPER or AAV-GFP was injected into C57BL/6 mice and then fed them HFD for eight weeks. (**a**) Food intake in AAV-GFP or AAV-BMPER (B) injected mice that were fed HFD or CC diet. (**b**) Body weight. (**c-d**) Fed insulin and glucose. (**e-f**) Glucose and insulin tolerance tests. n=6 (**a**), 8 (**b-f**). *, *P*< 0.05. **, *P*< 0.01. ****, *P*< 0.0001. NS, not significant.

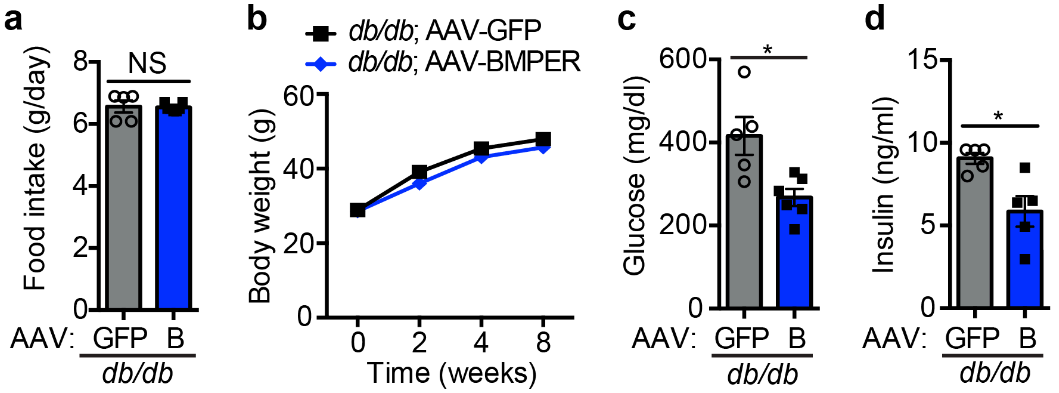


**Supplementary Figure 8. AAV-BMPER improves glucose responses in *db/db* mice.** AAV-BMPER or AAV-GFP was injected into 5-week-old *db/db* mice and then monitored for eight weeks. (**a**) Food intake in AAV-GFP or AAV-BMPER (B)- injected *db/db* mice. (**b**) Body weight. (**c-d**) Fed insulin and glucose. n=5 for AAV-GFP and 7 for AAV-BMPER group. *, *P*< 0.05. NS, not significant.
